# Cancerous phenotypes associated with hypoxia-inducible factors are not influenced by the volatile anesthetic isoflurane in renal cell carcinoma: Hypoxia-inducible factors are not activated by isoflurane

**DOI:** 10.1101/375311

**Authors:** Chisato Sumi, Yoshiyuki Matsuo, Munenori Kusunoki, Tomohiro Shoji, Takeo Uba, Teppei Iwai, Hidemasa Bono, Kiichi Hirota

## Abstract

The possibility that anesthesia during cancer surgery may affect cancer recurrence, metastasis, and patient prognosis has become one of the most important topics of interest in cancer treatment. For example, the volatile anesthetic isoflurane was reported in several studies to induce hypoxia-inducible factors, and thereby enhance malignant phenotypes *in vitro*. Indeed, these transcription factors are considered critical regulators of cancer-related hallmarks, including “sustained proliferative signaling, evasion of growth suppressors, resistance to cell death, replicative immortality, angiogenesis, invasion, and metastasis.” This study aimed to investigate the impact of isoflurane on the growth and migration of derivatives of the renal cell line RCC4. We indicated that isoflurane treatment did not positively influence cancer cell phenotypes, and that hypoxia-inducible factors (HIFs) maintain hallmark cancer cell phenotypes including gene expressions signature, metabolism, cell proliferation and cell motility. The present results indicate that HIF activity is not influenced by the volatile anesthetic isoflurane.

## Introduction

The hypothesis that anesthesia during cancer surgery may affect tumor recurrence, metastasis, and patient prognosis [1, 2] is gaining currently increasing importance [3]. Accordingly, numerous recent *in vitro*, *in vivo*, retrospective, and translational studies have investigated the the effect of anesthetics on perioperative immunity and cancer metastatic potential. For example, isoflurane was reported in several studies to induce hypoxia-inducible transcription factors (HIFs), and thereby enhance malignant phenotypes *in vitro* [4–6]. HIF-1 was originally cloned as a driver of erythropoietin expression [7–10]; however, shortly thereafter, it was reportedly associated with the tumor grade in various cancers [11]. Indeed, HIFs are now well-known as critical regulators of cancer hallmarks, including “sustained proliferative signaling, evasion of growth suppressors, resistance to cell death, replicative immortality, angiogenesis, invasion, and metastasis” [12, 13]. Moreover, tumor suppressors such as TP53 and PTEN regulate HIFs. Another striking example of the physiological significance of HIFs is von Hippel-Lindau (VHL) disease, a hereditary cancer syndrome predisposing individuals to highly angiogenic tumors, wherein the constitutive overexpression of vascular endothelial growth factor and glucose transporter 1 can be rectified corrected by functional VHL protein, a tumor suppressor that targets HIFs for degradation. This study aimed to investigate the effect of the volatile anesthetic isoflurane on growth and migration of derivatives of the renal cell line RCC4 that express (RCC-VHL) or do not express (RCC4-EV) VHL [14]. The present results indicate that HIFs significantly influence cancer cell growth and migration; however, isoflurane does not affect HIF-dependent phenotypes.

## Materials and methods

### Cell culture and reagents

Renal cell carcinoma cell lines stably transfected with pcDNA3-VHL (RCC4-VHL) or empty pcDNA3 (RCC4-EV) were kindly provided by Dr. Hiroshi Harada (Kyoto University) [15]. These cells were maintained in Dulbecco’s modified Eagle’s medium supplemented with 10% fetal bovine serum, 100 U/mL penicillin, and 0.1 mg/mL streptomycin. Purified mouse antibodies to human HIF-1α (Clone 54/HIF-1α) were purchased from BD Biosciences (San Jose, CA), while rabbit monoclonal antibodies againt HIF-1β/ARNT (D28F3) XP were purchased from Cell Signaling Technology (Danvers, MA). Antibodies againt HIF-2 α/EPAS1 were obtained from Novus Biologicals (Littleton, CO). Isoflurane and mouse monoclonal antibodies to α-tubulin were obtained from FUJIFILM Wako Pure Chemical (Osaka, Japan) [15–17] (Table 1).

**Table 1.**
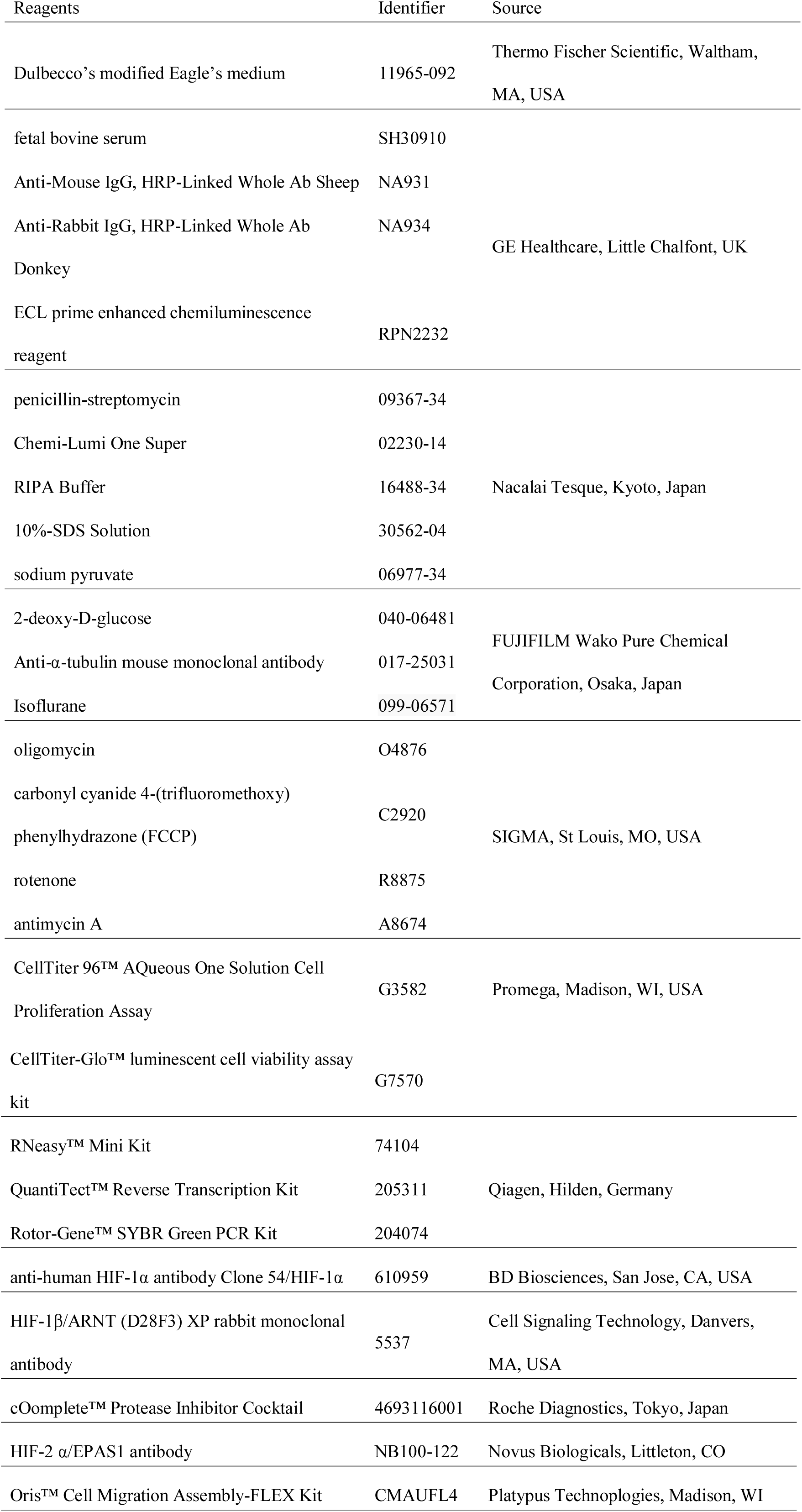
Key Resources Table.

### Induction of hypoxia and isoflurane treatment

Cells were maintained in an airtight chamber or workstation (AS-600P; AsOne, Osaka, Japan) perfused with air (MODEL RK120XM series; Kofloc, Kyotanabe, Japan) mixed with or without test anesthetic delivered by a specialized vaporizer within the open circuit. Gases and anesthetics were monitored during the experiment on an anesthetic gas monitor (Type 1304; Bluël & Kjær, Nærum, Denmark) calibrated with a commercial standard gas of 47% O_2_, 5.6% CO_2_, 47% N_2_O, and 2.05% sulfur hexafluoride [18, 19].

### Immunoblotting

Whole-cell lysates were prepared by incubating cells for 30 min in cold radioimmune precipitation assay buffer with cOmplete™ Protease Inhibitor Cocktail Tablets (Roche Diagnostics, Tokyo, Japan). Samples were then centrifuged at 10,000 ×g to sediment the cell debris, and 35 μg total protein from the resulting supernatant was resolved via 7.5% sodium dodecyl sulfate-polyacrylamide, gel electrophoresis and electro-transferred to membranes, probed with 1:2,000 of the indicated primary antibodies, probed with 1:8,000 of donkey anti-rabbit IgG (GE Healthcare, Piscataway, NJ) or sheep anti-mouse IgG (GE Healthcare) conjugated with horseradish peroxidase, and visualized with enhanced Chemi-Lumi One Super (Nacalai Tesque, Kyoto, Japan) (S1 Figure). Bands were quantified via densitometric analysis using Image Studio Lite (LI-COR Biosciences, Lincoln, NE, USA) [15, 20, 21]. Experiments were performed in triplicate, and representative blots are shown. Detailed protocols are available at protocols.io (dx.doi.org/10.17504/protocols.io.x9mfr46).

### Semi-quantitative real-time reverse transcriptase-polymerase chain reaction analysis (*q*RT-PCR) analysis

Total RNA was extracted from cells using RNeasy™ Mini Kit (Qiagen, Hilden, Germany) according to the manufacturer’s instructions. First-strand synthesis and RT-PCR were performed using QuantiTect™ Reverse Transcription Kit (Qiagen) and Rotor-Gene™ SYBR Green PCR Kit (Qiagen), following the manufacturer’s protocol. Targets were amplified and quantified in Rotor-Gene Q™ (Qiagen) and the expression fold-change of each target mRNA was normalized to that of β-actin. SLC2A1 was quantified with the QuantiTect Primer Assay QT00068957, while VEGFA was quantified with forward primer 5’-AGCCTTGCCTTGCTGCTCT-3’ and reverse primer 5’-TCCTTCTGCCATGGGTGC-3’. HIF1A was quantified using the forward and reverse primers 5’-ACACACAGAAATGGCCTTGTGA-3’ and 5’-CCTGTGCAGTGCAATACCTTC-3’, while EPAS1(HIF2A) was quantified with the forward and reverse primers 5’-ATGGGACTTACACAGGTGGAG-3’ and 5’-GCTCTGTGGACATGTCTTTGC-3’.

### Assessment of cell growth

Cell growth was assessed using Proliferation Assay™ Kit in accordance with the manufacturer’s instructions (Promega, Madison, WI, USA) [15, 21, 22]. Briefly, cells were seeded in 96-well plates (2 × 10^3^ cells/well), cultured overnight, treated with or without 2% isoflurane for 2 h, and then incubated in 5% CO_2_ and 95% air for indicated times. Cells were then incubated for 30 min at 37 °C with 20 μL of CellTiter 96 AQueous One Solution™, and the resulting absorbance at 490 nm was measured on an iMark™ microplate reader (Bio-Rad, Hercules, CA, USA). Cell viability was calculated relative to that of cells not treated with isoflurane, which was defined as 100%. All samples were tested in triplicate or quadruplicate per experiment. Detailed protocols are available at protocols.io (dx.doi.org/10.17504/protocols.io.v64e9gw).

### ATP assay

Intracellular ATP levels were quantified with CellTiter-Glo™ Luminescent Cell Viability Assay Kit (Promega, Madison, WI) in accordance with the manufacturer’s instructions [15, 22]. Briefly, cells were seeded in 96-well plates (2 × 10^3^ cells/well), cultured overnight, treated with or without 2% isoflurane for 2 h, incubated in 5% CO_2_ and 95% air for indicated times, reacted for 10 min with 100 μL CellTiter-Glo reagent, and assayed on an EnSpire^TM^ Multimode Plate Reader (PerkinElmer, Waltham, MA, USA). In this reaction, luminescence is generated by luciferase, which requires ATP to oxidize luciferin. Detailed protocols are available at protocols.io (dx.doi.org/10.17504/protocols.io.v7ke9kw).

### Cellular oxygen consumption and extracellular acidification rate

Cellular oxygen consumption rate and extracellular acidification rate were determined with XF Cell Mito Stress Test™ and XF Glycolysis Stress Test™, respectively, using an XFp Extracellular Flux Analyzer™ (Seahorse Bioscience, USA) [15, 21, 22]. RCC4-EV and RCC4-VHL cells (1 × 10^4^ cells/well) were seeded in an XFp cell culture microplate, cultured overnight, treated with or without 2% isoflurane for 2 h, and incubated in 5% CO_2_ and 95% air for 6 h. Oxygen consumption was then assessed in glucose-containing XF base medium in accordance with the manufacturer’s instructions, using a sensor cartridge hydrated at 37 °C in a non-CO2 incubator the day before use. Injection port A was loaded with 1.0 μM oligomycin (complex V inhibitor), port B was loaded with 2.0 μM carbonyl cyanide-4-(trifluoromethoxy) phenylhydrazone (FCCP), and port C was loaded with 0.5 μM rotenone/antimycin A (inhibitors of complex I and complex III). The sensor was calibrated using cells incubated at 37 °C in a non-CO2 incubator and in 180 μL of assay medium (XF base medium with 25 mM glucose, 1 mM pyruvate, and 2 mM L-glutamine, pH 7.4). The plate was immediately assayed after calibration. Assay parameters were calculated as follows: basal oxygen consumption rate = last rate before oligomycin injection − minimum rate after rotenone/antimycin-A injection; maximal oxygen consumption rate = maximum rate after FCCP injection − minimum rate after rotenone/antimycin A injection; non-mitochondrial oxygen consumption rate = minimum rate after rotenone/antimycin A injection; proton leak = minimum rate after oligomycin injection – non-mitochondrial respiration. To measure extracellular acidification rate, injection port A was loaded with 10 mM glucose, and the sensor was calibrated with cells incubated at 37 °C in a non-CO2 incubator and in 180 μL of assay medium (XF base medium with 2 mM L-glutamine, pH 7.4). The plate was immediately assayed after calibration and loading with oligomycin (1 μM) and 50 mM 2-deoxy-D-glucose. Extracellular acidification rate was normalized to total protein/well and calculated as extracellular acidification rate (glycolysis) = maximum rate after glucose injection − last rate before glucose injection. Detailed protocols are available at protocols.io (dx.doi.org/10.17504/protocols.io.v92e98e).

### Cell migration assay

Cell migration was analyzed using Oris™ Cell Migration Assay (Platypus Thechnologies, Madison, WI). Cells (2 × 10^4^ cells/well)) were seeded in wells plugged with stoppers to restrict seeding to outer areas only. Cells were then exposed for 8 h to 21% oxygen and 5% carbon dioxide balanced with nitrogen with or without 2% isoflurane. Stoppers were then removed to expose unseeded sites, into which cells could migrate during subsequent incubation at 37 °C in 5% CO_2_ and 95% air for indicated times. Cell migration was imaged on a BZ-9000 Fluorescence Microscope (KEYENCE, Itasca, IL), and non-colonized areas were quantified in pixels in ImageJ 1.51 (National Institutes of Health), corrected for total unseeded area, and expressed as percentage of colonized areas in reference wells.

### RNA sequencing

Total RNA was extracted from cells using RNeasy Mini Kit (Qiagen) [15, 21], and processed using TruSeq Stranded mRNA Sample Prep Kit (Illumina, San Diego, CA, USA). Poly(A) RNA libraries were then constructed using TruSeq Stranded mRNA Library Preparation Kit (Illumina, San Diego, CA, USA), and sequenced at 100 bp paired-ends on an Illumina HiSeq 2500 platform. Sequencing data in FASTQ format were deposited in the DDBJ Sequence Read Archive under accession numbers DRR100656 (RCC4-EV cells), DRR100657 (RCC4-VHL cells), DRR111123 (RCC4-EV cells treated with isoflurane), and DRR111124 (RCC4-VHL cells treated with isoflurane).

FASTQ files were evaluated as described previously [15, 21]. In brief, gene lists for Metascape analysis were generated from the Cuffdiff output file of significantly differentially expressed genes (*p* < 0.05; S1 Table). Gene ontology annotations were extracted in Ensembl Biomart [23], and sorted by the common logarithms of ([FPKM of RCC4-EV] + 1) / ([FPKM of RCC4-VHL] + 1), which were calculated from the same Cuffdiff output file (S1 Table). We added 1 to FPKM values because it is not possible to calculate the logarithm of 0. Histograms were generated in TIBCO Spotfire Desktop v7.6.0 with the “Better World” program license (TIBCO Spotfire, Palo Alto, CA, USA). Detailed protocols are available at protocols.io (dx.doi.org/10.17504/protocols.io.x9qfr5w).

### Statistical analysis

Experiments were repeated at least twice with triplicates of each sample. Data are mean ± SD. Groups were compared in Prism 7™ (GraphPad Software, Inc. La Jolla, CA) by one-way analysis of variance or Dunnett’s test for post hoc analysis. *P*-values less than 0.05 indicated statistical significance.

## Results

### HIF-1 and HIF-2 expression and activity in RCC4-EV and RCC4-VHL cells

Initially, we investigated the protein expression of both HIF-1 and HIF-2. Two-hour exposure of isoflurane reportedly increased both HIF-1α and HIF-2α protein expression in RCC4 cells lacking VHL expression within 6 h [24]. We adopted two different protocols for isoflurane treatment. HIF-1α and HIF-2α protein were detected at 20% O_2_ in RCC4-EV cells, but not in RCC4-VHL cells (Fig 1A). Similar levels of expression were observed in the former at 1% O_2_ (Fig 1A). In addition, expression in RCC4-EV cells was insensitive to 2% isoflurane at 20% O_2_ (Fig 1B). In contrast, HIF-1α and HIF-2α were barely detectable in RCC4-VHL cells even after exposure to 20% O_2_ for 8 h with or without 2% isoflurane (Fig 1B). However, HIF-1α and HIF-2α were induced in RCC4-VHL cells at 1% O_2_ for 8 h (Fig 1A), but were suppressed by 2% isoflurane (Fig 1C). To investigate possible protocol-dependent effects, HIF-1α was also quantified in RCC4-EV and RCC4-VHL cells exposed to isoflurane for 2 h and then incubated for 6 h at either 20% or 1% O_2_ (Fig 1D). In RCC4-EV cells, HIF-1α was insensitive to isoflurane regardless of subsequent exposure to 20% or 1% O_2_, but was suppressed in RCC4-VHL cells subsequently exposed to 1% O_2_. Similar trends were observed in cells exposed to isoflurane for 8 h at 20% and 1% O_2_ (Figs 1E and 1F). Nonetheless, HIF-1β expression was stable in both cells regardless of isoflurane treatment.

**Fig 1.**
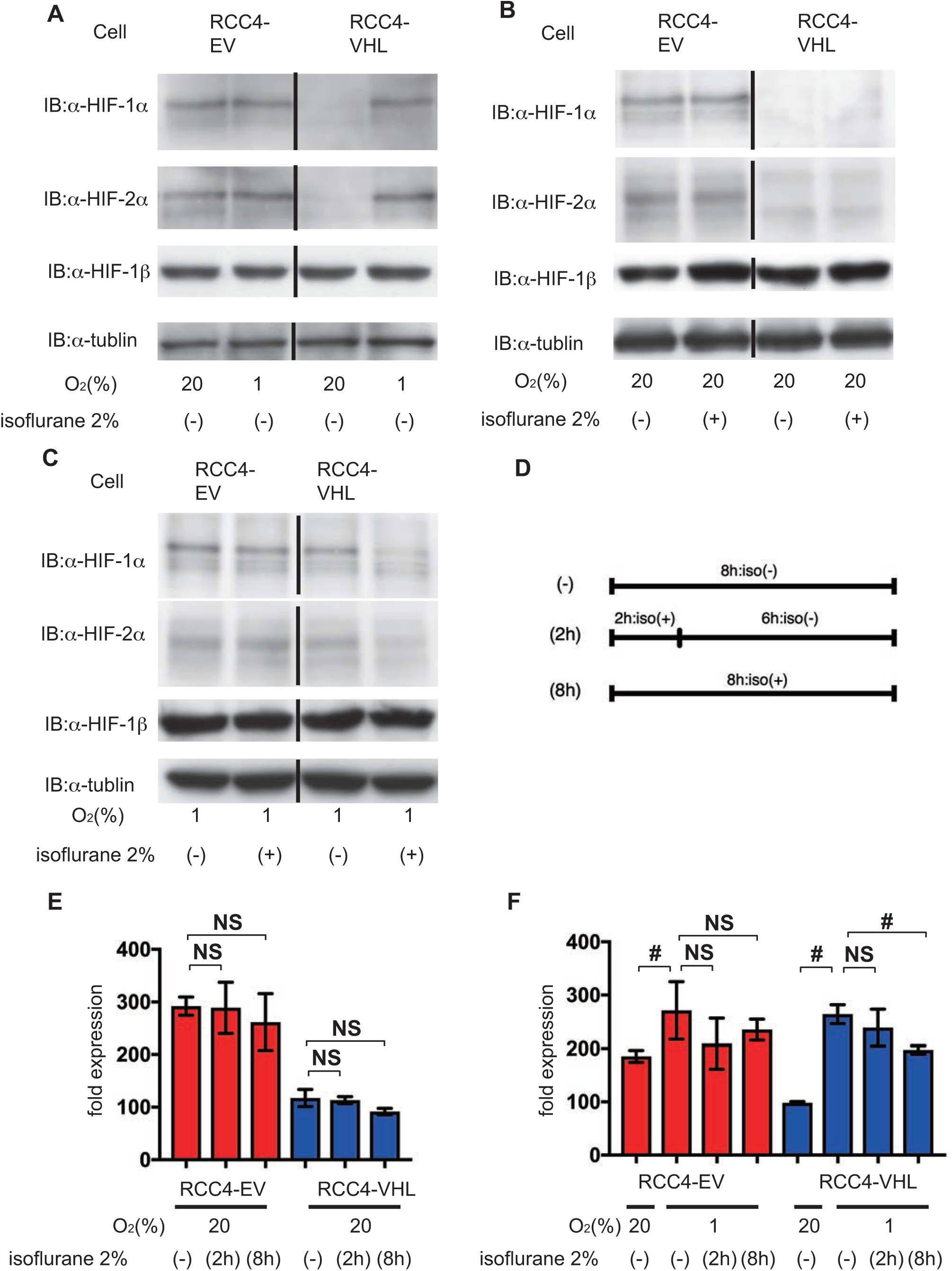
Expression of HIFs under isoflurane. (A) HIF expression after 8 h at 20% and 1% O_2_, (B) after 8 h at 20% O_2_ with or without 2% isoflurane, (C) and after 8 h at 1% O_2_ with or without 2% isoflurane. Cells were harvested, and 35 μg whole-cell lysates were immunoblotted using primary antibodies for the indicated proteins. (D) A schematic representation of two protocols of isoflurane treatment. (E and F) Cells were exposed to 2% isoflurane by two different protocols as indicated. HIF-1α expression was analyzed by densitometry and normalized to that in RCC4-VHL cells at 20% O_2_, which was considered 100%. Data represent the mean ± SD value; #, *p* < 0.05; NS, not significant.

Furthermore, we investigated expression of the HIFs-α subunits including HIF-1α and HIF-2α and HIF-downstream genes such as glucose transporter 1(*SLC2A1*) and vascular endothelial growth factor A *(VEGFA*) by semi-quantitative RT-PCR. *SLC2A1* and *VEGFA* were more abundant in RCC4-EV cells than in RCC4-VHL cells, but were induced in the latter at 1% O_2_ (Figs 2A and 2B). However, expression in RCC4-VHL cells at 1% O_2_ was suppressed by isoflurane. Interestingly, *HIF1A* and *EPAS1* (HIF-2α) mRNAs were less abundant in RCC4-EV cells, but were insensitive to isoflurane (Figs 2C and 2D). These results show that two different protocols for isoflurane treatment did not activate HIF-1 or HIF-2 under 20% O_2_ conditions.

**Fig 2.**
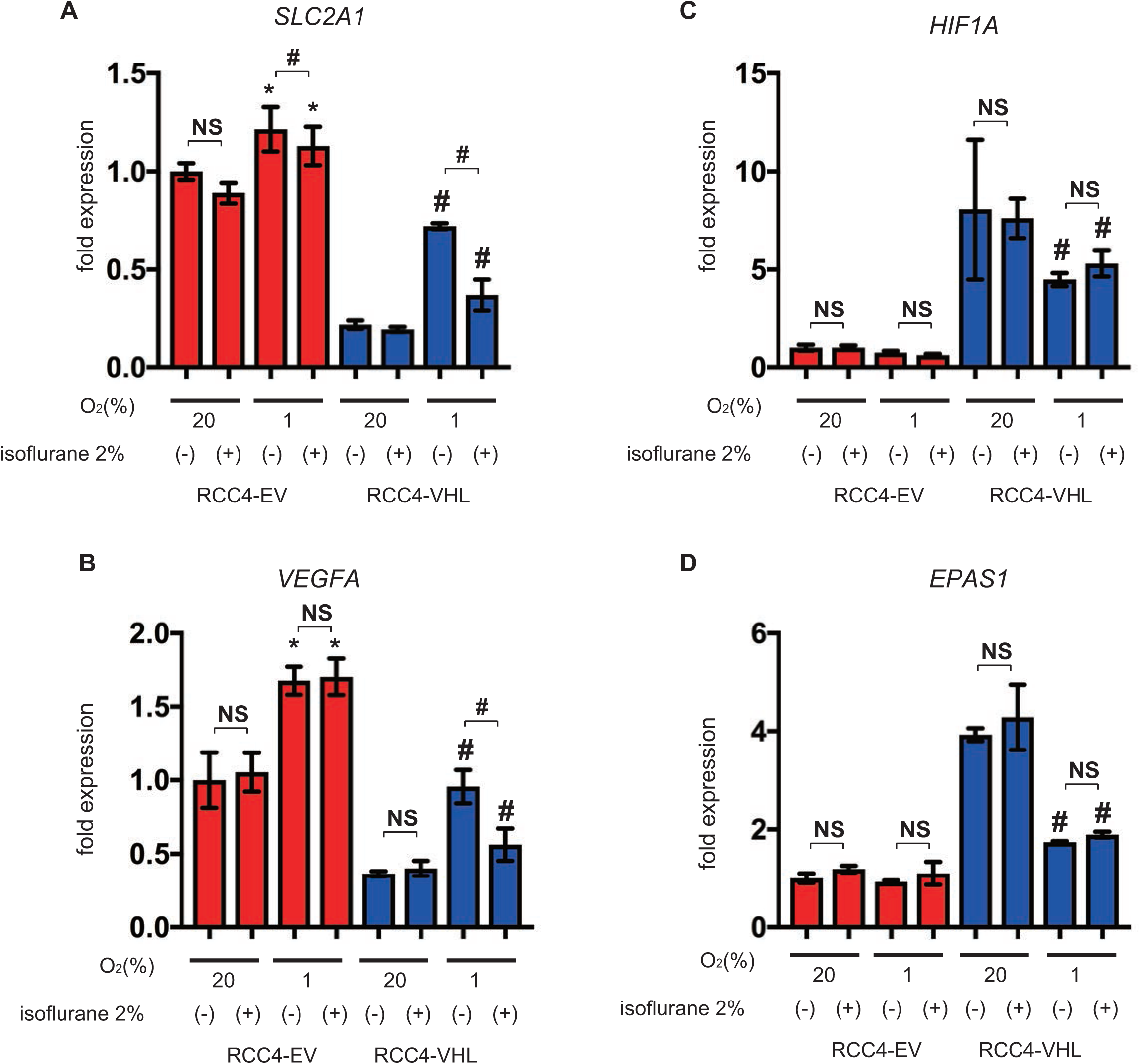
Expression of HIF-1 target genes under isoflurane. (A-D) RCC4-EV and RCC4-VHL cells were exposed for 8 h to 20% or 1% O_2_ with or without 2% isoflurane. Cells were then harvested, and mRNA levels quantified by semi-quantitative RT-PCR analysis. Relative expression fold-changes were determined from mRNA expression in RCC4-EV cells at 20% O_2_. Data represent the mean ± SD values. *, *p* < 0.05 vs. cells at 20% O_2_ and no isoflurane; #, *p* < 0.05 for the indicated comparison; NS, not significant; *SLC2A1*, solute carrier family 2 member 1; *VEGFA*, vascular endothelial growth factor A; *HIF1A*, hypoxia-inducible factor 1 α subunit; *EPAS1*, endothelial PAS domain protein 1.

### Effect of isoflurane on cell growth

Next, the critical phenotype of cancer cell cell growth was investigated. MTS assay is based on the reduction of MTS tetrazolium compound by viable cells to generate a colored formazan product that is soluble in cell culture media. This conversion is considered to be carried out by NAD(P)H-dependent dehydrogenase enzymes in metabolically active cells. Thus, the assay is correlate with cell growth. Cell growth was higher in RCC4-EV cells than in RCC4-VHL cells, but was insensitive to isoflurane (Fig 3A). Cell proliferation of RCC4-EV was suppressed by treatment with the HIF inhibitor YC-1 at 24 h (Fig. 3B). Similarly, cellular ATP was more abundant in the former than in the latter, and was also insensitive to isoflurane treatment (Fig 3C). In contrast, ATP levels were also decreased by treatment with YC-1 at 24 h (Fig. 3D). Our result strongly suggest that isoflurane treatment did not affect cell growth. However, HIFs are involved in cell growth and cellular energy metabolism.

**Fig 3.**
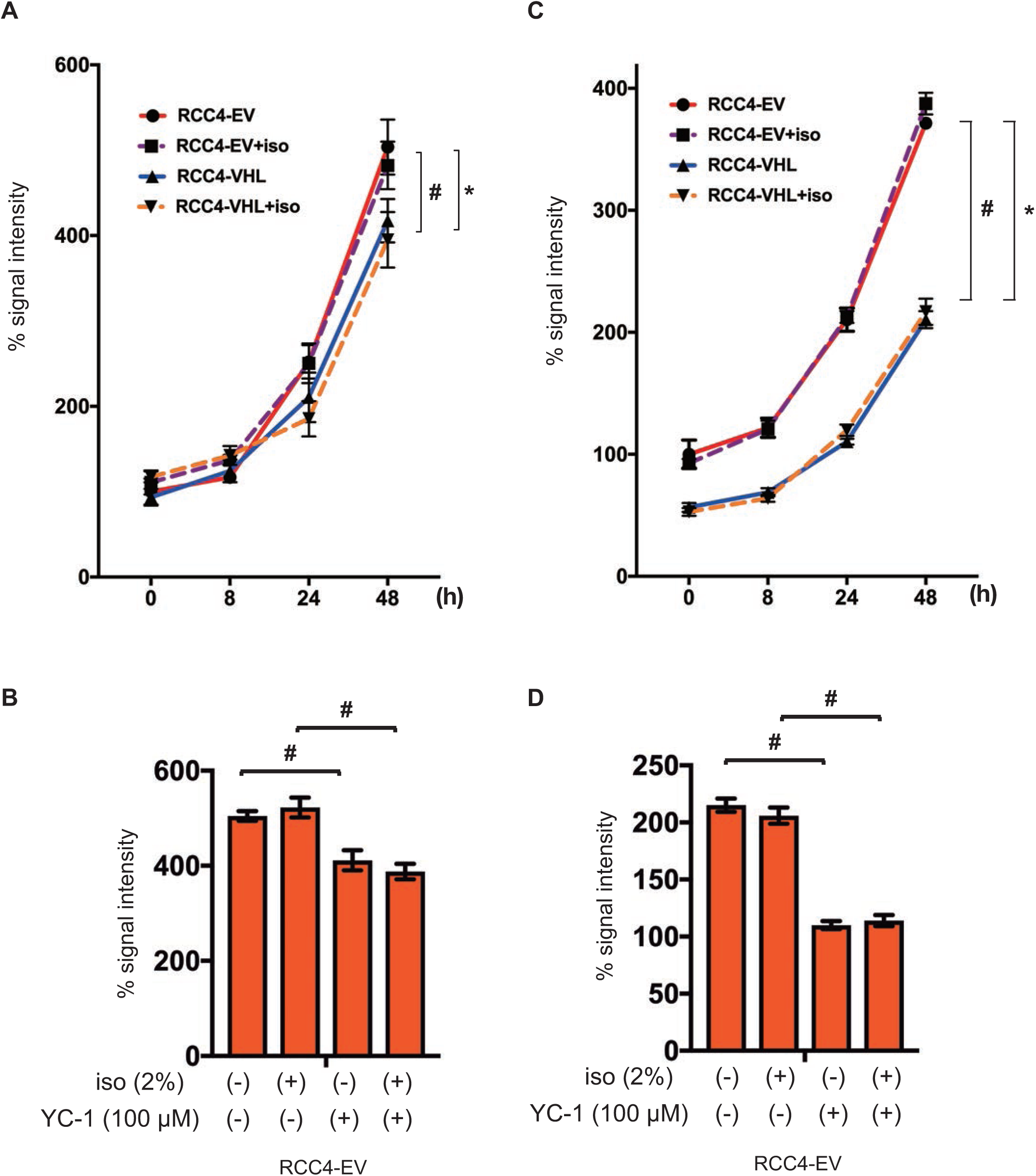
Cell proliferation under isoflurane. RCC4-EV and RCC4-VHL cells were grown for 48 h with or without exposure to 2% isoflurane for 2 h (A and C). RCC4-EV were grown for 24 h with or without exposure to 100 μM YC-1 or 2% isoflurane (B and C). (A and B) Cell growth at indicated time points, as evaluated by MTS [3-(4,5-dimethylthiazol-2-yl)-5-(3-carboxymethoxyphenyl)-2-(4-sulfophenyl)-2H-tetrazolium] assay. (C and D) Cellular ATP at indicated time points. Data represent the mean ± SD values. ^*^*p* < 0.05, for the comparison between RCC4-EV and RCC4-VHL cells with isoflurane treatment, #*p* < 0.05, for the comparison between RCC4-EV and RCC4-VHL cells without isoflurane treatment.

### Effect of isoflurane on cell migration

High cell motility is also one of the most significant feature of cancer cells. Therefore we examined the effect of isoflurane and HIFs on cell migration ability. RCC4-EV cells migrated significantly faster than RCC4-VHL cells over 12 h (Fig 4A), although exposure to 2% isoflurane for 2 h significantly suppressed migration in both cells (Fig 4B). The effect of isoflurane was concentration-dependent (S2 Figure). Furthermore, the involvement of HIF was examined. The HIF inhibitor YC-1 canceled HIF-dependent facilitation of cell migration but not the isoflurane-dependent suppression (Fig 4C). Similar to cell growth, cell motility depended on HIF activity but was not affected by isoflurane treatment.

**Fig 4.**
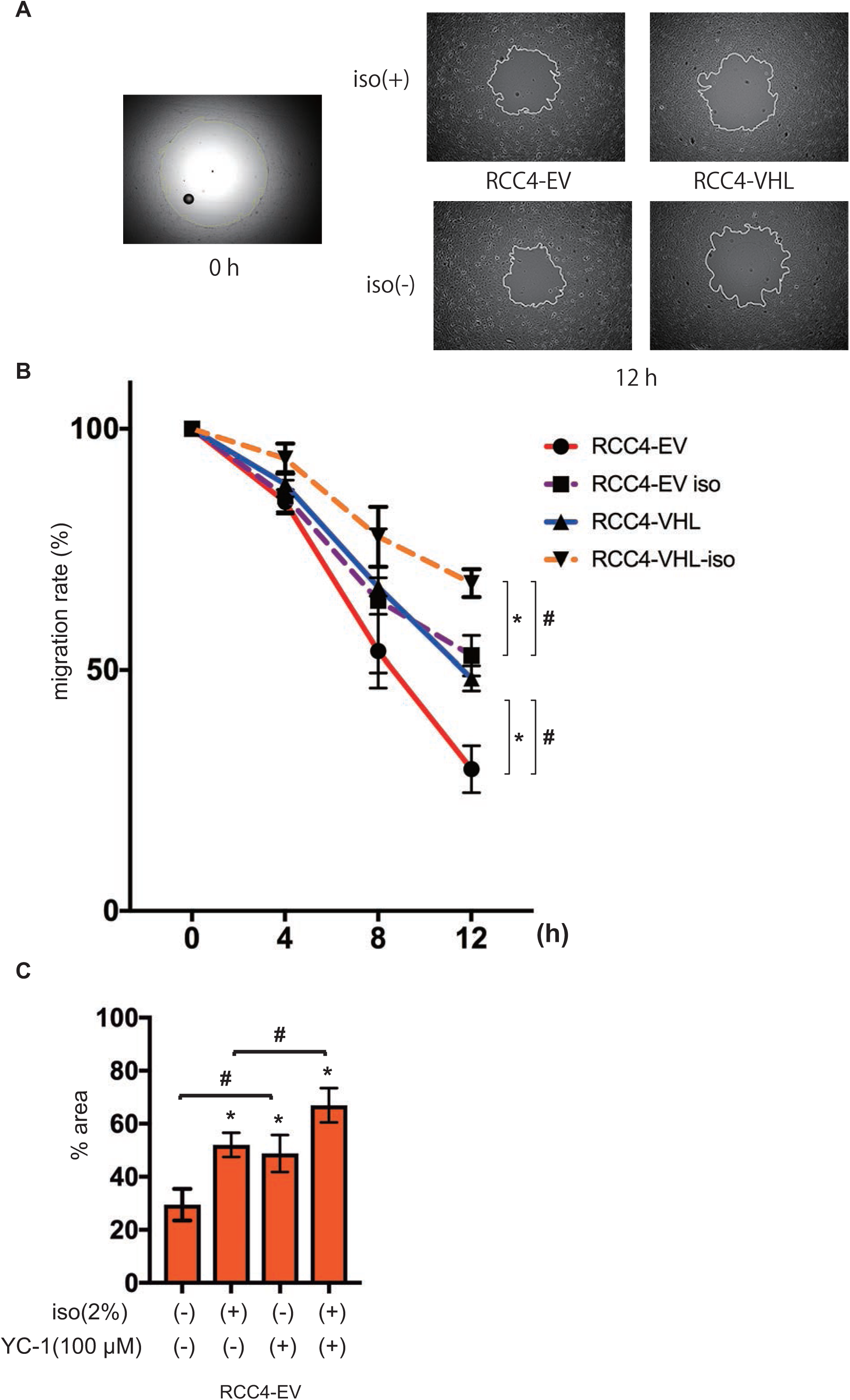
Cell migration under isoflurane. RCC4-EV and RCC4-VHL cells were allowed to migrate for 0–12 h after exposure 0% or 2% with or without 2% isoflurane treatment for 2 h. Migration rate was assessed as described in Materials and Methods. (A) Cell migration at 0 h and 12 h. (B) Non-colonized areas were quantified. “migration rate” is defined as (areas at each time / area at time 0) ×100. Data represent the mean ± SD values. ^*^*p* < 0.05, for the comparison between RCC4-EV and RCC4-VHL cells with or without isoflurane treatment, #*p* < 0.05, for the comparison between isoflurane (+) and isoflurane (-) groups in RCC4-EV and RCC4-VHL cells. (C) RCC4-EV cells treated with 100 μM YC-1 and allowed to migrate for 12 h with or without 2% isoflurane treatment for 2 h.

### Effect of isoflurane on glucose metabolism

In comparison to normal cells, cancer cells exhibit the Warburg effect, and thus preferentially metabolize glucose by glycolysis, producing lactate as an end product, despite availability of oxygen. Using an Extracellular Flux Analyzer™, the mitochondrial oxygen consumption rate was found to be lower in RCC4-EV cells in comparison to RCC4-VHL cells (Fig 5A), but was insensitive to isoflurane regardless of protocol (Fig 5B). However, extracellular acidification rate was higher in RCC4-EV cells relative to RCC4-VHL cells (Fig 5C), but was also insensitive to isoflurane (Fig 5D). Key parameters determining the mitochondrial oxygen consumption rate, including basal oxygen consumption rate, maximum respiration, proton leak, and nonmitochondrial respiration, were also calculated from Cell Mito Stress Test™ data (Fig 6). These parameters were significantly different between RCC4-EV and RCC4-VHL cells, but were insensitive to isoflurane.

**Fig 5.**
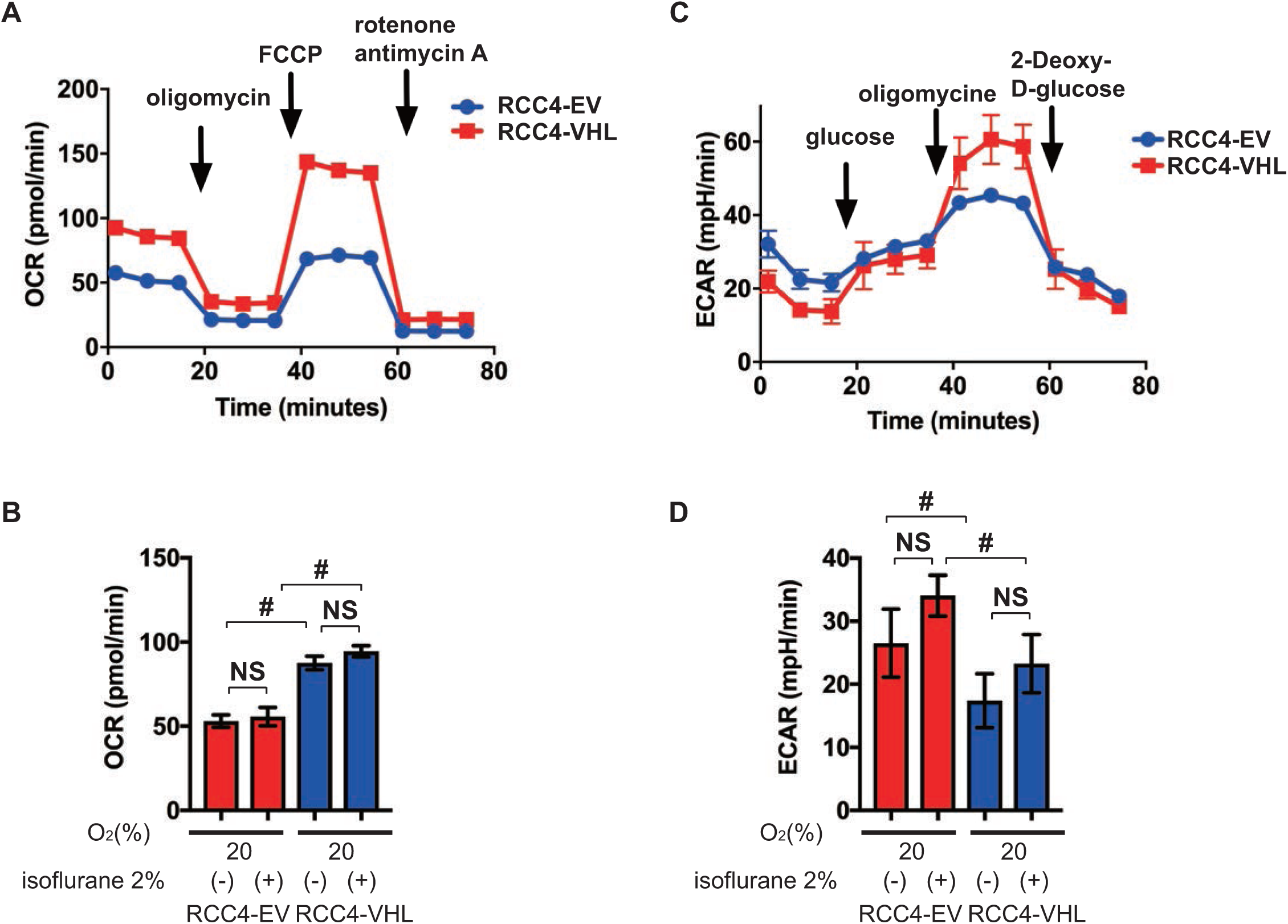
Oxygen metabolism in RCC4-EV and RCC4-VHL cells. (A) Cell Mito Stress Test™ at 20% O_2_. (B) Oxygen consumption rate with or without 2% isoflurane treatment for 2 h. (C) Glycolysis test at 20% O_2_. (D) Extracellular acidification rate with or without 2% isoflurane treatment for 2 h. #, *p* < 0.05 for the indicated comparison.

**Fig 6.**
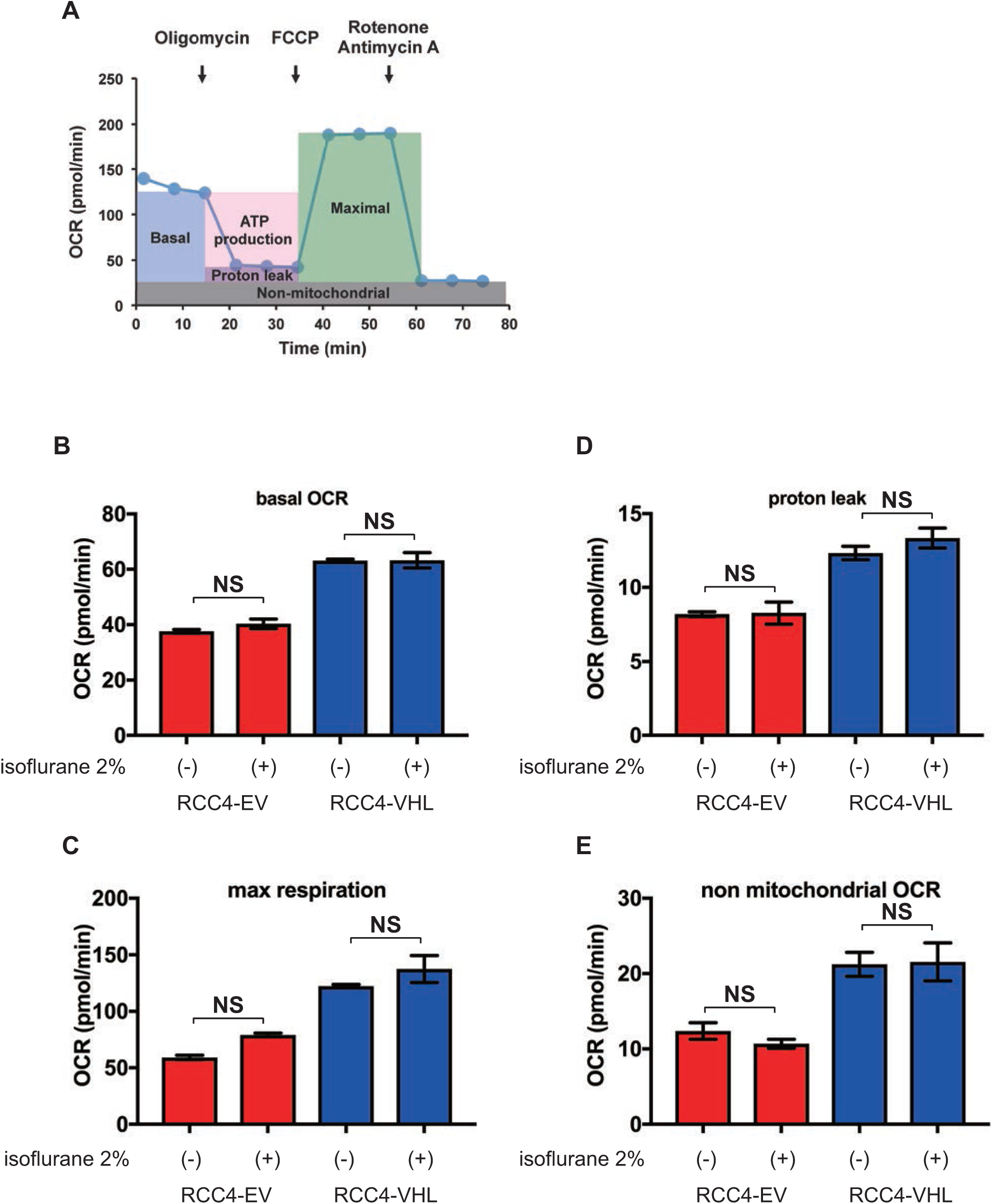
Reprogrammed oxygen metabolism in RCC4-EV and RCC4-VHL cells. (A) Cell Mito Stress Test™ profile of key parameters that determine mitochondrial oxygen consumption rate. (B) Basal oxygen consumption rate, (C) maximal oxygen consumption rate, (D) proton leakage, and (E) nonmitochondrial respiration rate with or without 2% isoflurane for 2 h.

### Effect of isoflurane on global gene expression

Persistent effects of isoflurane on the cancer cell phenotypes would result in significant changes in the gene expression landscape. Genome-wide gene expression patterns were assessed via RNA-Seq, using next-generation sequencing.

Clustering of RNA sequencing data (S1 Table) indicated that transcriptomic bias due to isoflurane was smaller than transcriptomic variations due to VHL expression (Fig 7A). Indeed, more than 200 genes were differentially expressed between RCC4-EV and RCC4-VHL cells, as inferred from Wilcoxon signed rank test of FPKM values at significance level 0.05 (Fig 7B). However, only one gene was differentially expressed in RCC4-VHL cells exposed to isoflurane, while no such gene was identified in RCC4-EV cells. Pairwise scatter plots comparing log_10_[FPKM+1] values from four experiments further confirmed this result (Fig 7C). Furthermore, enrichment analysis also revealed that GO:0001666 (response to hypoxia), GO:0010035 (response to inorganic substance), hsa05230 (central carbon metabolism in cancer), GO:003198 (extracellular matrix organization), and GO:0097190 (apoptotic signaling pathway) were significantly enriched in RCC4-EV cells (Fig 7D) regardless of isoflurane exposure (Fig 7E). Finally, only 42 genes annotated to cancer hallmark gene ontologies were sensitive to isoflurane, although the effects were negligible (Fig 8). Indeed, only CITED1 was strongly responsive to isoflurane.

**Fig 7.**
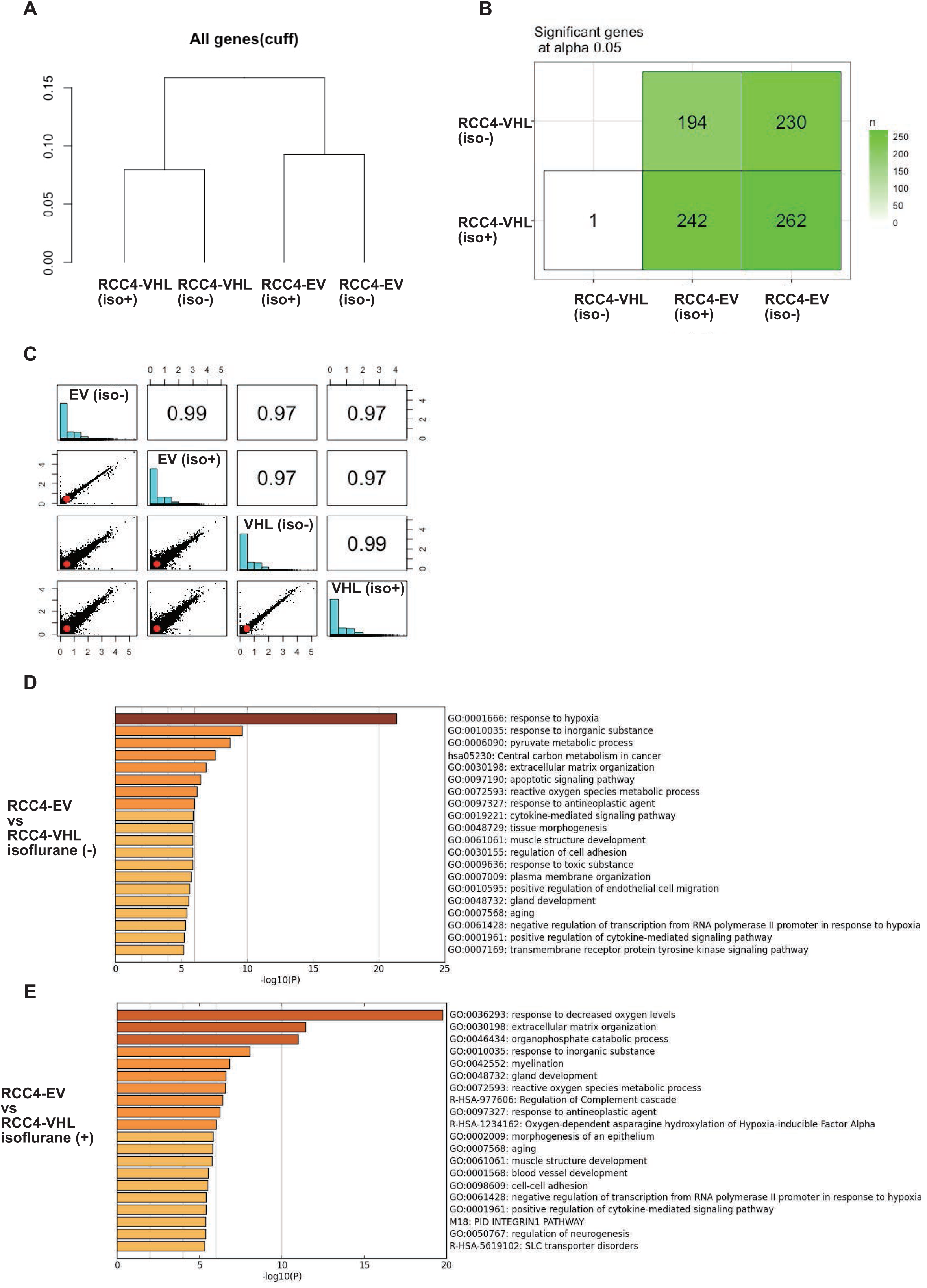
Global gene expression in RCC4-EV and RCC4-VHL cells. (A) Hierarchical clustering and (B) number of differentially expressed genes identified by pairwise Wilcoxon signed rank test at *p* < 0.05. (C) Scatter plots below the diagonal are pairwise comparisons of log_10_[FPKM+1]. Histograms at the diagonal show the distribution of genes by level of expression, and numbers above the diagonal are pairwise correlation coefficients. (D and E) Heatmap of enriched terms in differentially expressed genes, colored by *p* values and as inferred by Metascape analysis (http://metascape.org/) using the Cuffdiff output file (gene_exp.diff).

**Fig 8.**
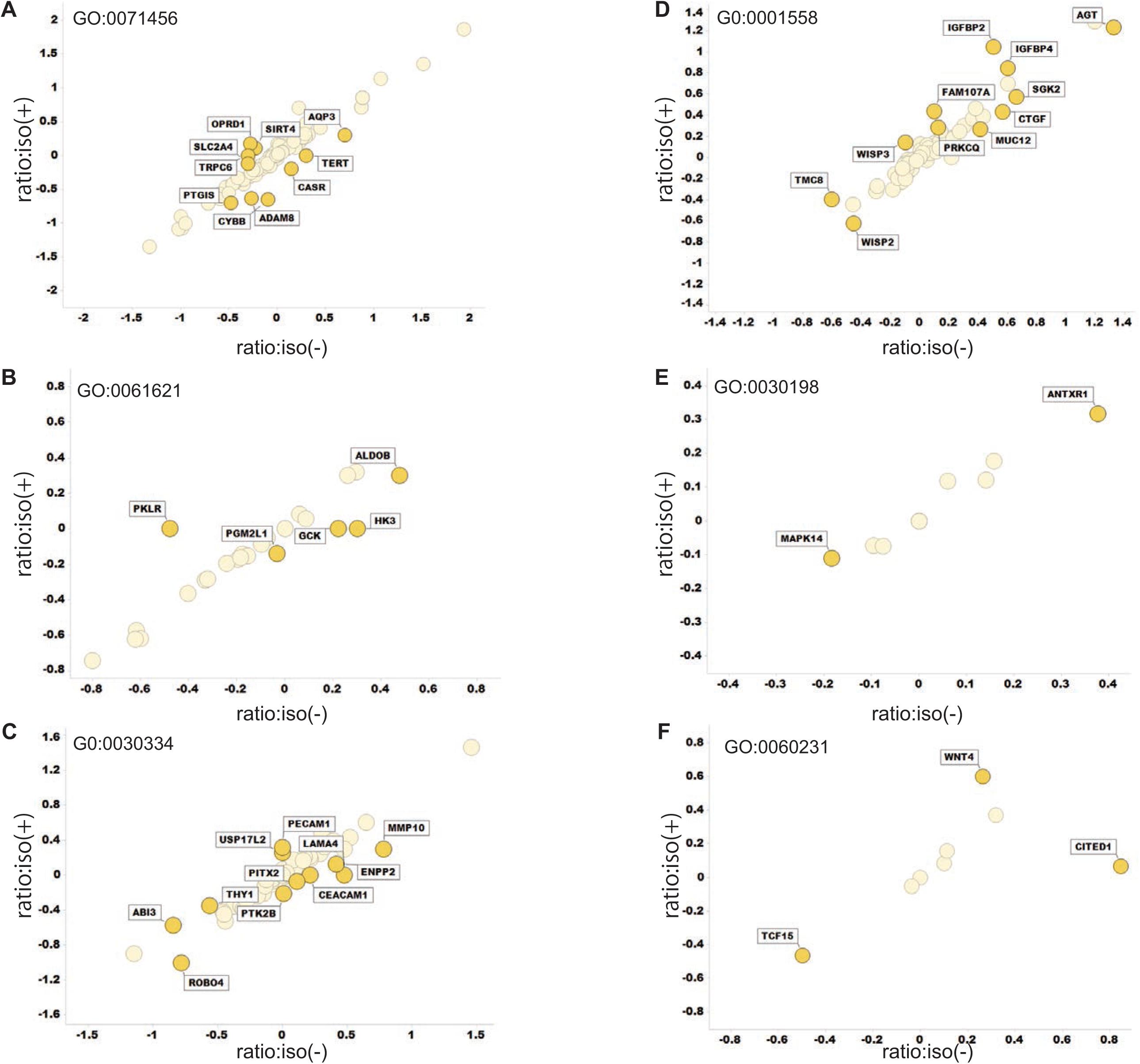
Transcriptomic variations in gene ontologies related to cancer hallmarks. Logarithms of ([FPKM of RCC4-EV] + 1)/([FPKM of RCC4-VHL] + 1) were plotted for cells treated with or without isoflurane on the Y and X axis, respectively. Data were calculated from the same Cuffdiff output file described in Fig 7. (A) GO:0071456 cellular response to hypoxia, (B) GO: 0061621 canonical glycolysis, (C) GO: 0030334 regulation of cell migration, (D) GO: 0001558 regulation of cell growth, (E) GO: 0030198 extracellular matrix organization, and (F) GO: 0060231 mesenchymal to epithelial transition.

## Discussion

This study shows that transcription factor HIFs maintain hallmarks of cancer cell phenotypes including gene expression signatures, metabolism, cell proliferation, and cell motility, and HIF activity is not influenced by the volatile anesthetic isoflurane. The hallmarks of cancer were originally proposed by Hanahan and Weinberg in 2000 [12], and included sustained proliferative signaling, evasion of growth suppressors, resistance to cell death, replicative immortality, angiogenesis, invasion, and metastasis. Subsequent conceptual progress has revealed two additional emerging hallmarks, namely reprogramming of energy metabolism and immune evasion [13]. Accordingly, we investigated the impact of isoflurane and HIF on these hallmarks through global gene expression. Since HIF has been extensively investigated in the context of cancer biology [4, 11], we used RCC4-EV cells, which are derived from human renal cell carcinoma. As these cells are VHL-deficient, both HIF-1 and HIF-2 are activated even under normoxic conditions, but are suppressed upon forced expression of VHL, as in RCC4-VHL cells [14]. Accordingly, RCC4-EV cells proliferated and migrated faster than RCC4-VHL cells, and exhibited metabolic reprogramming from oxidative phosphorylation to glycolysis. These results clearly indicate that HIFs are critically involved in cancerous phenotypes, as previously reported.

However, isoflurane treatment did not upregulate HIF-1α and HIF-2α expression in RCC4-EV cells. In contrast, isoflurane suppressed HIF-1α expression in RCC4-VHL cells at 1% O_2_. Accordingly, isoflurane also suppressed *SLC2A1* and *VEGFA*, which are downstream of HIF-1, under hypoxic conditions, although this effect is minor to that of VHL expression. A precedent study described that 2h isoflurane exposure upregulated both HIF-1α and HIF-2α protein expression in a PTEN-Akt-dependent manner in RCC4 cells lacking VHL, which is equivalent to RCC4-EV cells in this study, expression within 6 h [24]. In the study, HIF-1α and HIF-2α proteins are barely expressed without isoflurane treatment in RCC4-EV cells. However, the results are concurrent with previous reports including the landmark study although the reason is unknown [14, 25–27]. Hence, we investigated potential protocol-dependent effects; however, we found that neither exposure to isoflurane for 8 h, nor for 2 h followed by culture for another 6 h, affected HIF-1 and HIF-2 in RCC4-EV cells.

In 2006, Exadaktylos et al. [2] proposed that anesthesia and analgesia during cancer surgery may affect tumor recurrence or metastasis, a hypothesis that was subsequently supported by several clinical studies [1]. Potential underlying mechanisms may include direct cellular effects, as well as indirect effects on patient immunity and on cancer metastasis. In this study, we focused on hallmarks of cancer phenotypes in relation to HIF activity. The effect of isoflurane on proliferation, cell growth and cell migration were investigated. The precent results clearly indicate that isoflurane treatment did not exert positive effects on cancer cells positively. In addition, metabolism and global gene expression appeared to be sensitive to HIFs but not to isoflurane. This study has several limitations. The present data are derived entirely from *in vitro* experiments in established cell lines; however, xenografts may be required to elucidate the impact of anesthetics on cancer progression *in vivo*. In addition, since surgical procedures are aimed at eliminating most of the cancer, there is the complication that the remaining cells may be in an environment containing damaged tissue and inflammation; anesthetics may alter ongoing cell-cell interactions in such an environment.

In summary, this study shows that demonstrated that isoflurane does not affect HIF activity in renal carcinoma cells, nor the expression of genes associated with cancer hallmarks. However, we confirmed that HIFs help maintain cancerous phenotypes.

## Acknowledgments

We thank Editage (www.editage.jp) for English language editing.

## Author contributions

Conceptualization: Chisato Sumi, Yoshiyuki Matsuo, Kiichi Hirota.

Data curation: Chisato Sumi, Hidemasa Bono, Kiichi Hirota.

Funding acquisition: Chisato Sumi, Hidemasa Bono, Kiichi Hirota, Kiichi Hirota.

Investigation: Chisato Sumi, Munenori Kusunoki, Tomohiro Shoji, Takeo Uba, Teppei Iwai.

Supervision: Kiichi Hirota.

Writing – original draft: Chisato Sumi, Kiichi Hirota.

Writing – review & editing: Chisato Sumi, Yoshiyuki Matsuo, Munenori Kusunoki, Tomohiro Shoji, Takeo Uba, Teppei Iwai, Hidemasa Bono, Kiichi Hirota

## Additional information

The datasets analyzed in the current study are available in the Supplementary Information and can be obtained from the corresponding author on reasonable request.

