## Supplementary figures and images for "Cancerous phenotypes associated with hypoxia-inducible factors are not influenced by the volatile anesthetic isoflurane in renal cell carcinoma: Hypoxia-inducible factors are not activated by isoflurane"

### Supplementary S1

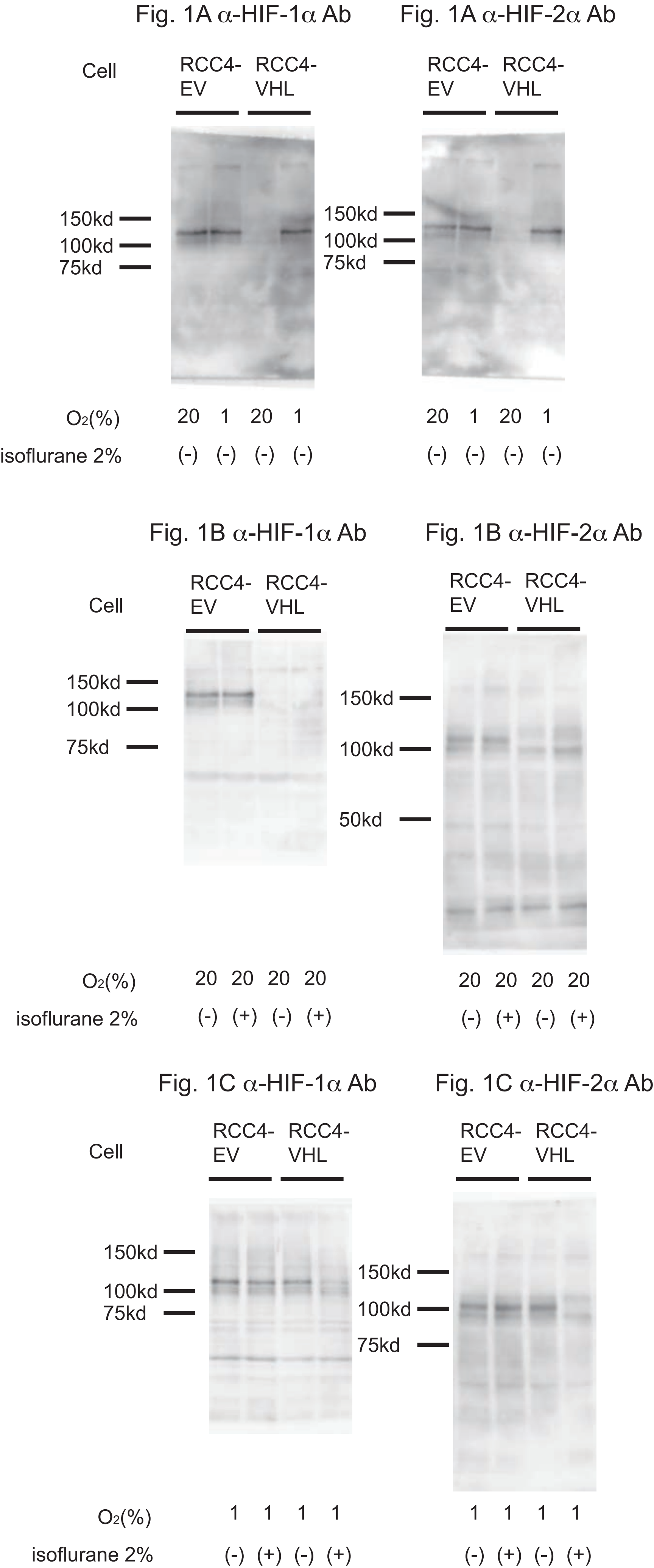

### Supplementary S2

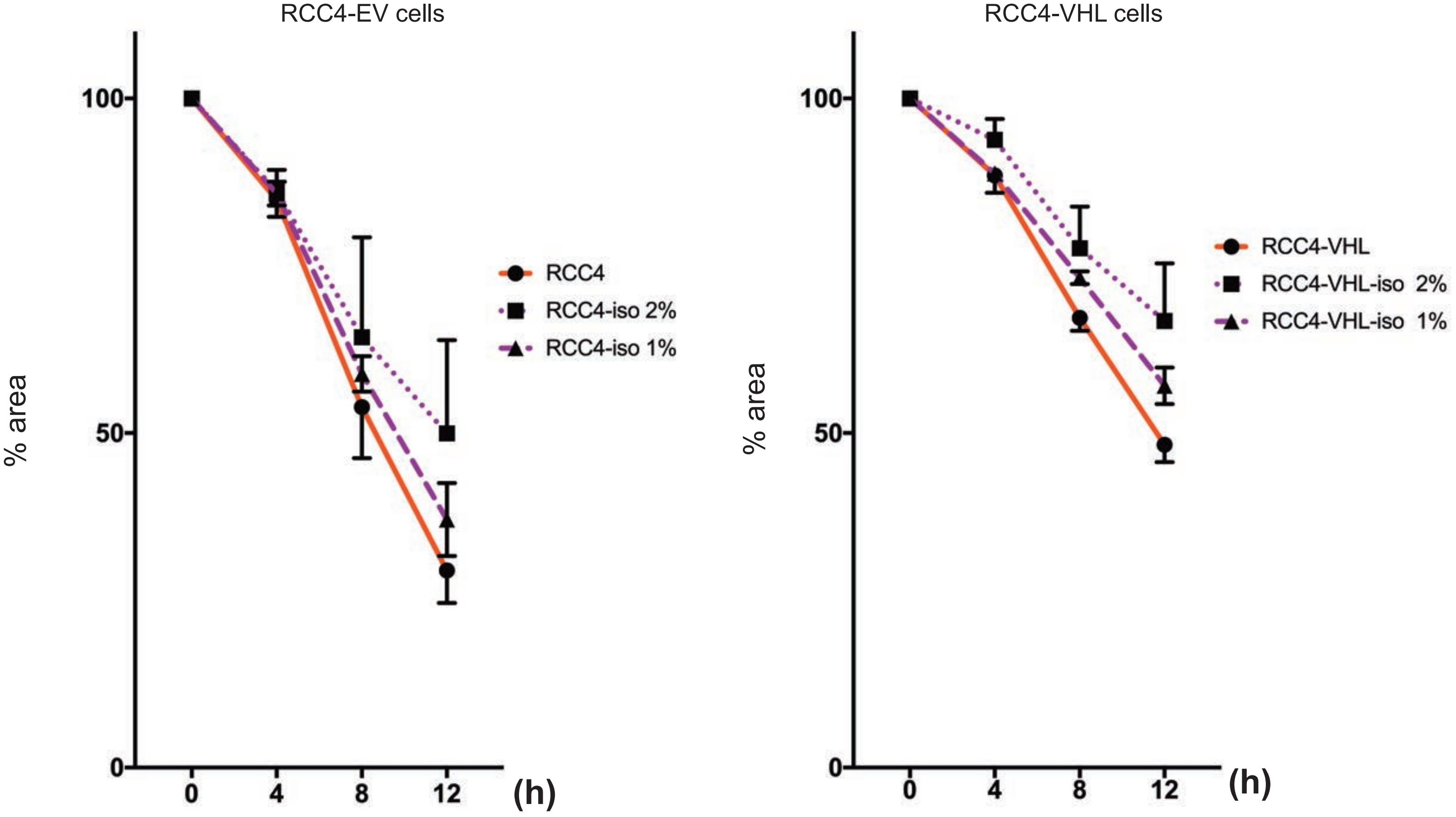
